# Intelligent systems distribute memory across internal and embodied resources

**DOI:** 10.64898/2026.10.16.759765

**Authors:** Ruiyi Zhang, Xaq Pitkow, Dora E Angelaki

**Affiliations:** Neuroscience Institute, Carnegie Mellon University; Machine Learning Department, Carnegie Mellon University; Tandon School of Engineering, New York University; Center for Neural Science, New York University

## Abstract

Intelligent systems do not operate in isolation from the body, yet how bodily actions support internal computation remains unclear. Here we show that primates and artificial agents recruit eye movements to maintain beliefs about hidden goals when internal computation is constrained. In a naturalistic navigation task, both systems spontaneously tracked goals with their eyes after they disappeared from view. This strategy emerged in artificial agents only when internal memory was constrained, identifying resource constraints as a driver of memory externalization. Restricting gaze impaired performance in both systems, while natural behavioral variability and causal perturbations in agents established a direct influence of oculomotor state on internal beliefs about hidden goals. Across agents, varying the reliability of internal memory and gaze redistributed goal information between internal and embodied states. The same resource-allocation principle predicted the distribution of goal information in macaque parietal cortex, where greater reliance on gaze was accompanied by weaker gaze-independent neural representations. Together, these findings identify eye movements as embodied memory and reveal a convergent principle across intelligent systems: memory is allocated across internal and embodied resources according to their reliability.

## Main

Intelligence emerges from an interplay between brain and body. Yet the two are typically treated asymmetrically: memory and inference are attributed to internal neural states, whereas bodily movements are viewed primarily as outputs of those computations [1, 2]. Internal computation, however, is necessarily limited: neural representations are noisy [3], capacity-limited [4, 5] and metabolically costly [6, 7]. Why, then, maintain information entirely within the brain when the body itself can carry some of the computational burden?

Theories of embodied cognition propose that actions can participate directly in computation rather than merely express its results [8, 9]. Humans count on their fingers [10], manipulate objects to simplify reasoning [8], and restructure their surroundings to reduce demands on memory and inference [11, 12]. Such behaviors show that bodily states and interactions can themselves serve as computational resources that reduce demands on internal computation, but leave a deeper question unresolved: what determines how computation is distributed across internal and embodied resources? One possibility is that resource reliability determines their relative use, with greater reliance on resources that provide more reliable task-relevant information. Whether such resource constraints govern how information is distributed across brain and body remains unclear.

Eye movements provide a tractable test of this hypothesis. When navigating towards a goal that is no longer visible, an animal must maintain an estimate of the goal location and update it continuously as it moves [13, 14]. Neural theories typically attribute this ability to internal states that encode and update spatial information over time [15, 16]. Yet primates also track invisible goals with their eyes during navigation [17, 18], raising the possibility that gaze itself carries part of the information required for memory. If so, gaze could provide an embodied memory resource for maintaining spatial information about hidden goals.

Here we use gaze to investigate whether resource constraints determine how memory is distributed between internal and embodied states. We first establish that primates and artificial agents independently recruit gaze to track hidden goals and that gaze contributes to memory-guided navigation. By manipulating internal memory and oculomotor reliability in artificial agents, we then identify the conditions under which this strategy emerges and establish how resource reliability governs memory allocation. Finally, we test whether the resulting resource-allocation principle predicts the organization of goal information in macaque parietal cortex. Together, we show that intelligent systems allocate memory across internal and embodied resources according to their reliability.

### Gaze behavior converges in primates and agents

Three rhesus macaques were trained to navigate through a virtual environment using a joystick (Fig. 1a, Methods). On each trial, a single target was randomly sampled from a range of possible locations and presented for 0.3 s before disappearing (Fig. 1b), requiring the animals to maintain and continuously update an estimate of the hidden goal during navigation. The environment contained no landmarks that could specify the hidden target location. Throughout each trial, subjects controlled their linear and angular velocity using the joystick and were free to move their eyes (Fig. 1c). Navigation therefore engaged two independently controlled motor systems: joystick actions changed the animal’s location in the environment, whereas oculomotor actions changed the state of the eyes (Fig. 1d). Subjects were rewarded only for stopping within a predefined boundary around the target.

**Figure 1:**
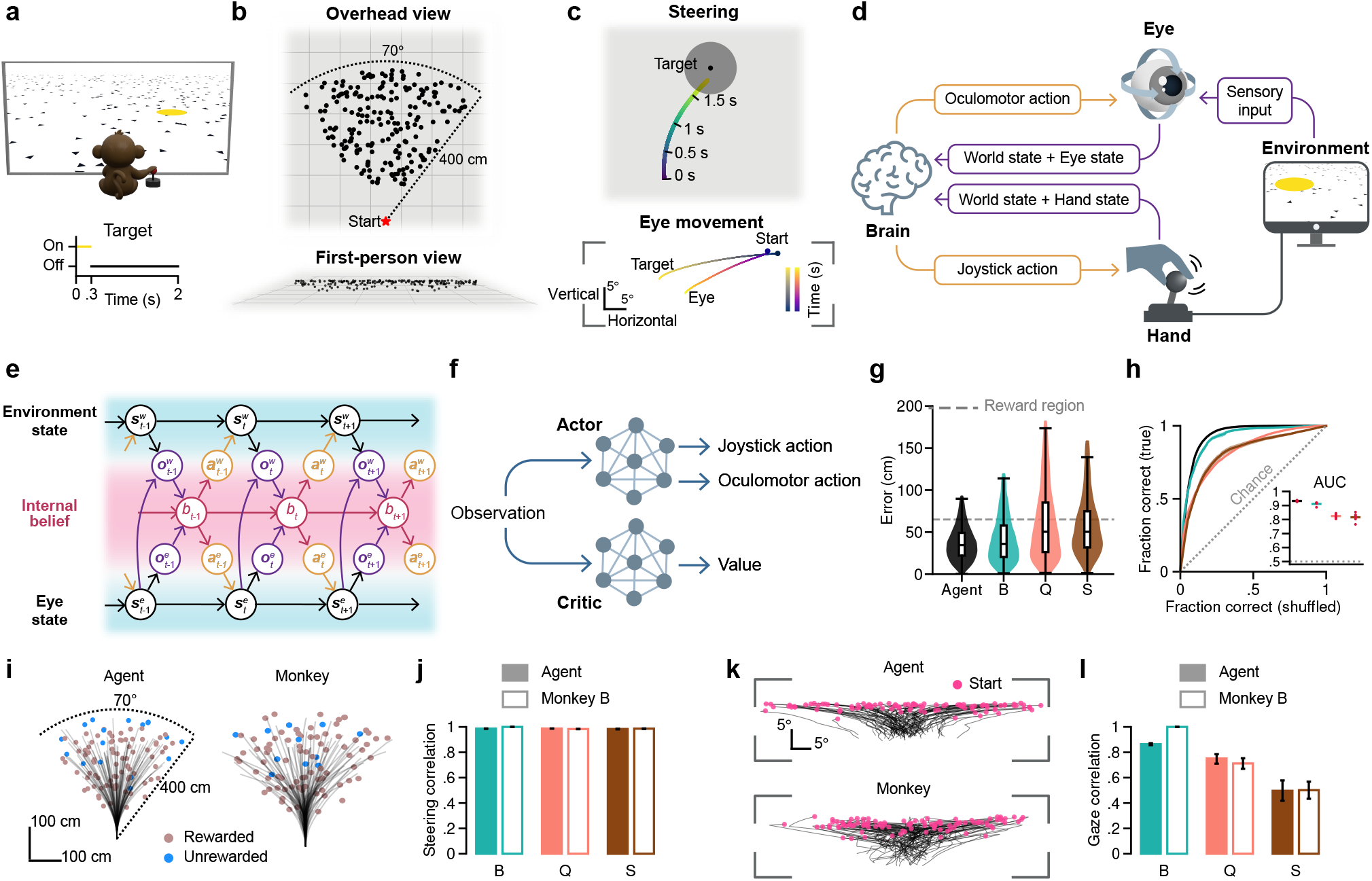
Gaze behavior converges in monkeys and artificial agents during memory-guided navigation. **a**, Virtual-reality navigation task. Subjects used a joystick to steer towards a briefly presented target, which disappeared after 0.3 s. **b**, Distribution of target locations (dots) shown from overhead (top) and first-person (bottom) views. **c**, Example trial showing the steering trajectory in world coordinates (top) and the corresponding eye and target trajectories in angular coordinates (bottom; Methods). **d**, Schematic of the sensorimotor loop. The brain generates oculomotor and joystick actions that shape bodily interactions with the environment, while sensory and bodily-state signals provide feedback to the brain. **e**, Graphical model of agent–environment interaction. Environment (***s***^*w*^) and eye (***s***^*e*^) states evolve through actions (***a***^*w*^, ***a***^*e*^), generating observations (***o***^*w*^, ***o***^*e*^) that update an internal belief (*b*) guiding subsequent actions. Eye state also shapes sensory observations of the environment. **f**, Actor–critic architecture of the artificial agent. Observations are processed by an actor that generates joystick and oculomotor actions and by a critic that estimates value. **g**, Distribution of navigation error for the agent and monkeys B, Q and S, measured as the Euclidean distance between target and final stopping location. Dashed line indicates the reward boundary. Embedded boxplots show quartiles. **h**, Receiver operating characteristic (ROC) analysis of navigation performance comparing stopping accuracy for true and shuffled target-response pairings across hypothetical reward boundaries (from Extended Data Fig. 1a; Methods). Dashed line denotes chance; inset shows area under the curve (AUC). Red dots denote individual training seeds (agent) or sessions (monkeys). **i**, Representative steering trajectories across target locations for an example agent and Monkey B. **j**, Similarity (Frobenius correlation) of time-aligned steering trajectories between each monkey and the agent (filled) or monkey B (open). Error bars indicate 95% confidence intervals (CI). **k**,**l**, As in **i**,**j**, but for eye trajectories. **g–l**, Analyses used 2,000 target-matched trials per subject, sampled across monkey sessions or six trained agent seeds.

We formalized this sensorimotor interaction as a dynamical system in which internal belief, environmental state and eye state evolve jointly through observation and action (Fig. 1e; Methods). Actions alter the environment and eyes, while resulting environmental and eye-state observations update internal belief; eye state can also shape sensory observations of the environment. We explicitly represented eye state because, unlike the joystick state that specifies locomotor commands, it provides an independently controllable state that can vary without determining the navigation trajectory. We instantiated this framework in recurrent reinforcement-learning (RL) agents [19] trained to perform the same navigation task (Fig. 1f; Methods). Network recurrence provided an internal state for temporal integration, while the actor generated joystick and oculomotor actions and the critic estimated value to guide learning. As in monkeys, reward depended only on navigation success; agents received no explicit objective or reward for their oculomotor behavior.

Both monkeys and agents navigated accurately towards remembered targets (Fig. 1g,h, Extended Data Fig. 1a). Their steering trajectories exhibited similar spatial structure across target locations, with agent–monkey correspondence approaching that observed between monkeys (Fig. 1i,j; Extended Data Fig. 1b; Methods). Strikingly, this convergence extended to eye movements: agents developed structured oculomotor dynamics that closely resembled those of the monkeys, despite receiving no explicit instruction or reward for gaze behavior (Fig. 1k,l and Extended Data Fig. 1c). Thus, biological and artificial systems independently converged on similar gaze behavior while solving the same memory-guided navigation problem.

### Gaze supports memory of hidden goals

The convergence of gaze behavior in monkeys and agents raised the possibility that eye movements carry information about the hidden goal. After target disappearance, gaze remained strongly correlated with target location in all three monkeys and the agent (Fig. 2a,b; Extended Data Fig. 2a–c; Methods). As navigation progressed, gaze became progressively more closely related to the eventual stopping location than to the true target (Extended Data Fig. 2d), suggesting that gaze reflected the subject’s evolving estimate of the hidden goal, including deviations from its true location.

**Figure 2:**
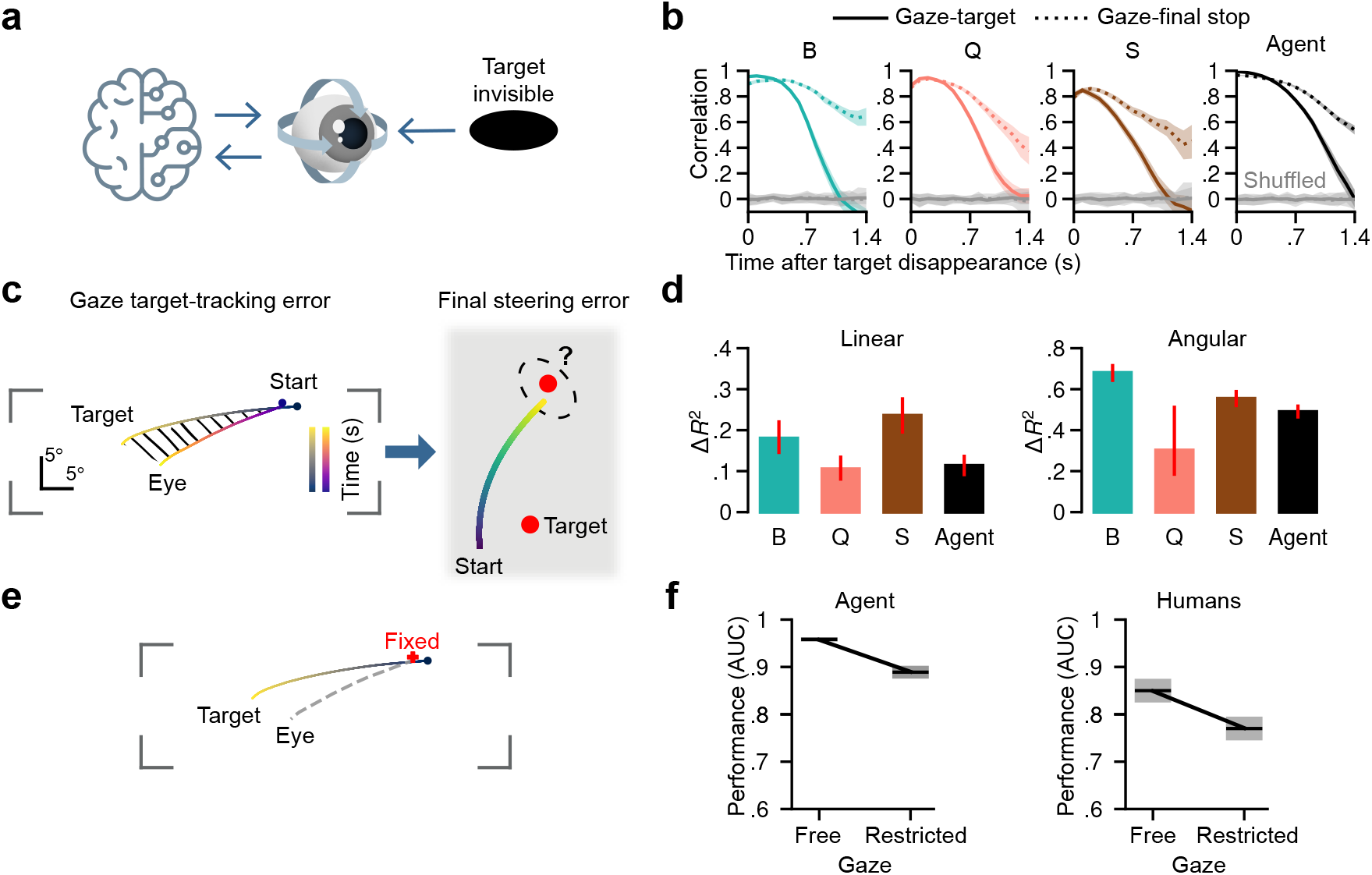
Gaze supports memory of hidden goals. **a**, Schematic of the relationship between gaze and hidden-target location. **b**, Frobenius correlation between gaze and angular target location over time after target disappearance for monkeys and the agent. Solid lines show gaze–target correlations; dotted lines show gaze–final-stop correlations. Grey lines indicate chance correlations obtained by shuffling gaze across trials. Shading indicates 95% CI. **c**, Schematic relating gaze target-tracking error during navigation to final steering error. Tracking error was quantified from gaze–target differences in angular coordinates, using Euclidean distance for linear steering error and signed horizontal difference for angular steering error. **d**, Additional variance in final linear (left) and angular (right) steering errors explained by gaze target-tracking error beyond target location (Δ*R*^2^), evaluated out of sample.

We therefore asked whether natural variability in gaze reflected trial-to-trial variability in the hidden-goal estimate. If so, gaze target-tracking errors should predict errors in the eventual stopping location (Fig. 2c). Because navigation difficulty varies with target location, we first accounted for target location and then asked whether gaze error explained additional trial-to-trial variation in steering error. Adding gaze error improved prediction of both linear and angular steering errors across monkeys and the agent (Fig. 2d; Extended Data Fig. 2e; Methods). Thus, errors in where subjects looked during navigation predicted errors in where they ultimately navigated.

To test the functional contribution of gaze to navigation performance, we prevented it from updating during subsequent navigation (Fig. 2e; Methods). In agents, eye position was frozen once gaze reached the visible target; in humans, participants were instructed to maintain gaze at the target’s last visible location on screen. Restricting gaze impaired navigation in agents, mirroring the impairment previously observed in humans performing the same task (Fig. 2f; [17]). Together, these results show that gaze carries behaviorally relevant information about the evolving hidden-goal estimate and contributes to memory-guided navigation.

Linear error denotes Euclidean distance between target and final stopping location; angular error denotes their angular difference. Error bars indicate 95% CI. **e**, Gaze-restriction manipulation. After reaching the visible target, eye position was held fixed in agents; human participants were instructed to maintain gaze at the target’s last visible location. Red dot indicates the fixed gaze location; grey dashed line illustrates the eye trajectory under free gaze. **f**, Navigation performance with free or restricted gaze in agents and humans. Performance was quantified by AUC as in Fig. 1h. Error bars indicate standard deviation (s.d.) across six agent seeds or human participants (*n* = 5); human data from [17]. **b**,**d**, Analyses used 2,000 target-matched trials per subject, sampled across monkey sessions or six trained agent seeds.

### Internal constraints drive memory externalization in agents

The contribution of gaze to memory raised a deeper question: what determines how strongly a system relies on gaze to maintain task-relevant information? We hypothesized that belief can draw on both internal and embodied resources (Fig. 3a), such that reliance on gaze should increase when internal resources become less reliable. The agents examined thus far had constrained internal memory; we therefore systematically varied this resource by adding different levels of noise to the actor’s recurrent state used for action selection (Methods), with greater noise reducing the reliability of internal memory. At each noise level, agents were trained with either free or restricted gaze. With reliable recurrent states, restricting gaze had little effect on navigation; as recurrent-state noise increased, however, the cost of gaze restriction grew markedly (Fig. 3b; Extended Data Fig. 3a). Thus, gaze became behaviorally important specifically when internal resources were limited.

**Figure 3:**
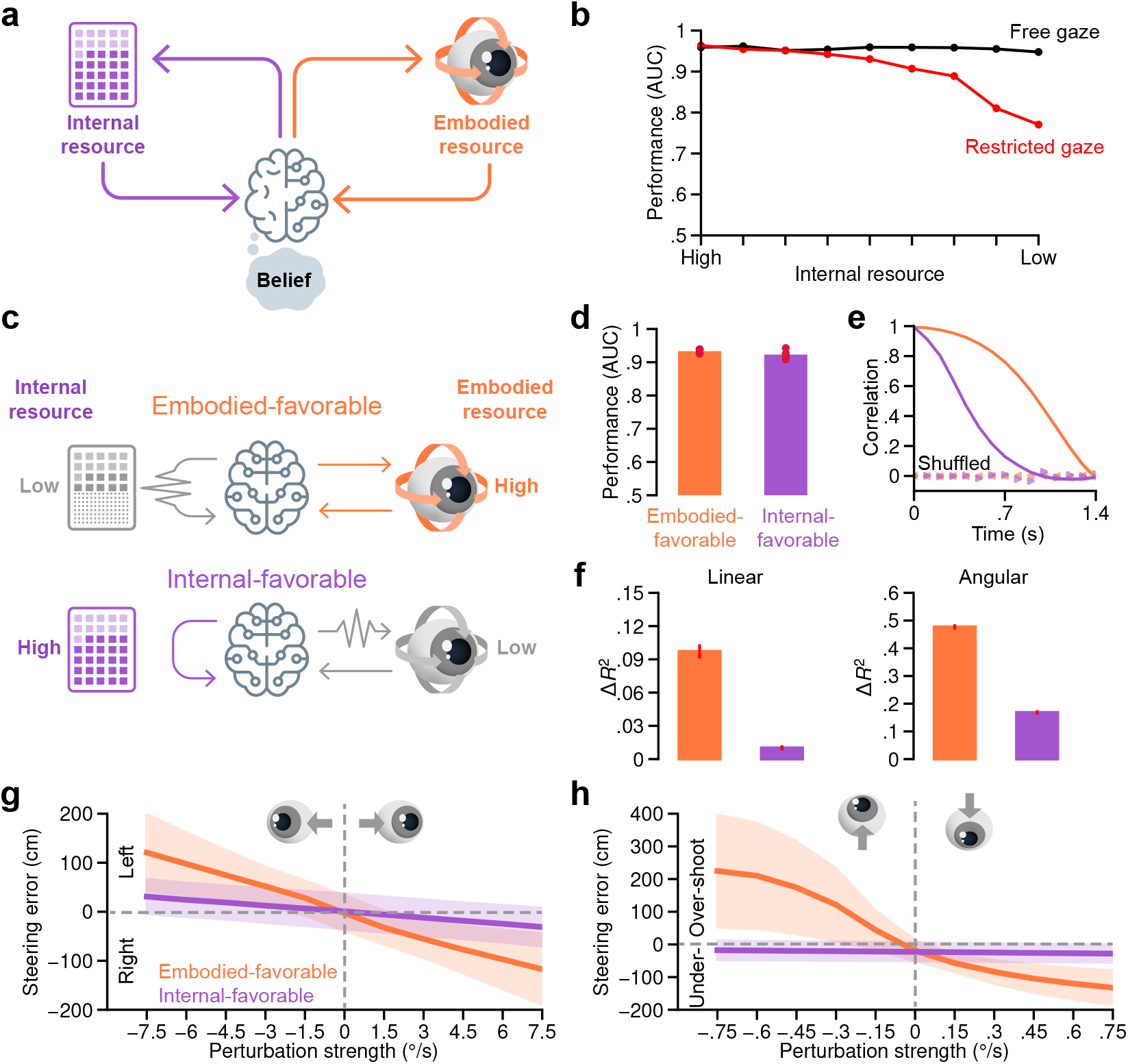
Internal constraints drive memory externalization. **a**, Hypothesis that belief can draw on internal and embodied resources. **b**, Navigation performance (AUC) of agents trained with free or restricted gaze as internal resources were progressively constrained by increasing recurrent-state noise. Points show means across six training seeds. Gaze restriction was applied as in Fig. 2e. **c**, Resource configurations of two agent conditions: an embodied-favorable condition with constrained internal memory and reliable oculomotor control, and an internal-favorable condition with reliable internal memory and noisy oculomotor control. **d**, Navigation performance (AUC) of embodied- and internal-favorable agents. Red dots denote individual training seeds. **e**, Gaze–target correlation over time for the two agent conditions, quantified as in Fig. 2b. Dots indicate correlations after shuffling gaze across trials; shading indicates 95% CI. **f**, Additional variance in final linear (left) and angular (right) steering errors explained by gaze target-tracking error beyond target location, as in Fig. 2d. Error bars indicate 95% CI. **g**,**h**, Effect of horizontal (g) and vertical (**h**) gaze perturbations introduced at test time in the two agent conditions. Positive and negative steering errors denote leftward and rightward errors in (**g**), and overshooting and undershooting in (**h**), respectively. Perturbations were applied for 1 s from the onset of navigation. Grey dashed lines indicate zero perturbation and zero steering error; shading indicates s.d. **e–h**, trials were pooled across six training seeds per agent condition.

We next asked whether changing the available resources produced distinct computational strategies. We compared two agent conditions with opposing resource configurations: an embodied-favorable condition combining constrained internal memory with reliable oculomotor control, and an internal-favorable condition combining reliable internal memory with noisy oculomotor control (Fig. 3c; Methods). Despite closely matched navigation performance (Fig. 3d), embodied-favorable agents showed stronger gaze–target correlation and greater prediction of final steering errors from gaze than internal-favorable agents (Fig. 3e,f; Extended Data Fig. 3b).

To test whether this difference reflected a causal contribution of eye state to the hidden-goal estimate, we perturbed gaze at test time and measured the resulting change in final stopping location, the behavioral readout of that estimate. In embodied-favorable agents, gaze perturbations produced graded, direction-dependent stopping errors: horizontal perturbations shifted lateral stopping position, whereas vertical perturbations produced under- or overshooting (Fig. 3g,h; Methods). These effects were substantially weaker in internal-favorable agents. Together, these results show that constraints on internal resources promote reliance on gaze, producing an embodied memory strategy in which eye state exerts greater influence over the internal estimate of the hidden goal.

### Reliability governs memory allocation in agents

The contrasting strategies in Fig. 3 suggested that memory might be flexibly distributed according to the resources available to the agent. We therefore systematically varied the reliability of both resources, crossing eight levels of recurrent-state noise with eight levels of oculomotor-control noise to generate 64 training conditions (Fig. 4a; Methods). Agents achieved broadly similar navigation performance across this resource landscape (Fig. 4b).

**Figure 4:**
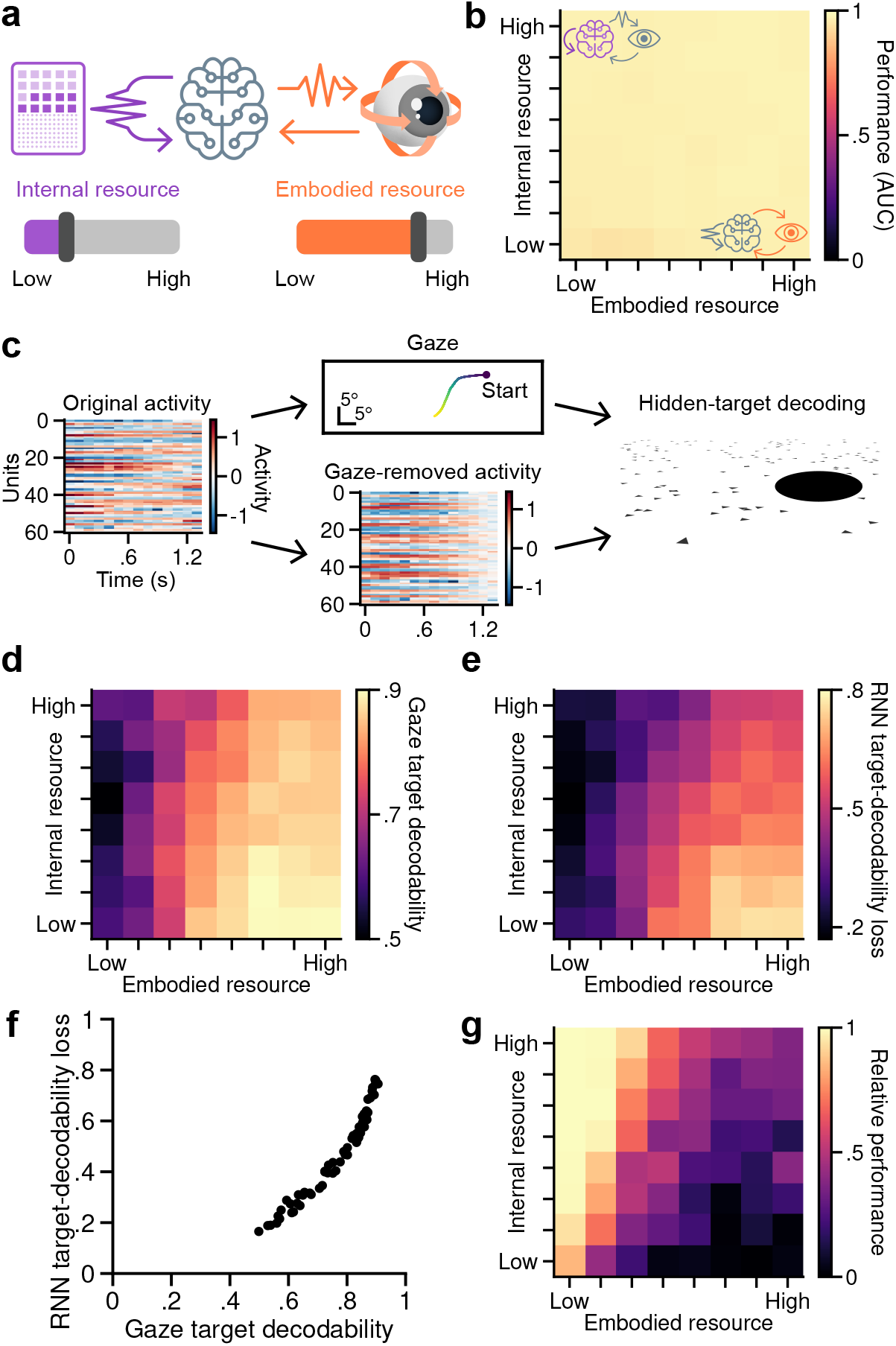
Reliability governs memory allocation across internal and embodied resources. **a**, Schematic illustrating independent variation of internal and embodied resources. **b**, Navigation performance (AUC) across 64 agent conditions spanning eight levels each of internal and embodied resources, manipulated through recurrent-state and oculomotor-control noise, respectively. **c**, Example trial after target disappearance illustrating the target-decoding analysis. Heatmaps show 60 example recurrent units over time before and after linear removal of gaze-related variance. Activity is shown as z-scores relative to the original activity of each unit. Hidden-target location in angular coordinates was decoded from gaze, original recurrent activity and gaze-removed recurrent activity. **d**, Decodability of hidden-target location from gaze across the resource landscape. **e**, Loss of target decodability from recurrent activity after removing gaze-related variance, defined as decodability from original minus gaze-removed activity. **f**, Relationship between gaze target decodability (**d**) and recurrent target-decodability loss (**e**) across the 64 resource conditions (Pearson’s *r* = 0.97, *P* = 5.1× 10^*−*39^). **d–f**, Decodability was quantified by Frobenius correlation between predicted and true target locations. **g**, Relative navigation performance under gaze restriction introduced at test time, defined as the fraction of rewarded trials with restricted gaze divided by that with free gaze. At each time step, oculomotor updating was blocked with probability 0.8. **b**,**d–g**, Values are averaged across eight training seeds per resource condition.

We next turned from behavior to the agents’ representations, asking how resource reliability shaped the target information carried by gaze and network activity. We used target decodability from gaze as a measure of embodied memory. Because network activity can reflect both gaze-related signals and information maintained independently of gaze, we removed gaze-related variance and decoded the target from the remaining activity as a measure of gaze-independent internal memory (Fig. 4c; Methods). As internal resources became less reliable and embodied resources more reliable, target information in gaze increased (Fig. 4d), while gaze-independent target information in the recurrent state decreased (Extended Data Fig. 4a,b), yielding a greater loss of RNN target decodability after gaze removal (Fig. 4e). Across resource conditions, target information in gaze closely tracked this RNN target-decodability loss (Fig. 4f). As a control, we created a null gaze signal by shifting gaze in time to disrupt its correspondence with network activity, then removed activity variance associated with this null signal before decoding. Target decodability remained close to its original level, showing that the reduction with true gaze depended on the gaze–activity relationship rather than removal procedure itself (Extended Data Fig. 4c,d). Together, these results show that the amount of hidden-goal information maintained in each resource increased with its relative reliability.

Finally, we tested whether this distribution of memory predicted functional reliance on gaze. We restricted oculomotor updating at test time across the same resource landscape and measured performance relative to free gaze (Fig. 4g). Gaze restriction had little effect when internal resources were reliable, but increasingly impaired performance as memory became more strongly embodied. Across resource conditions, greater allocation of target information to gaze predicted stronger impairment under gaze restriction (Extended Data Fig. 4e,f). Together, these results identify a resource-allocation principle: the relative reliability of internal and embodied resources governs how memory is distributed across them to guide behavior.

### Resource allocation predicts memory organization in the brain

We next asked whether the resource-allocation principle established in agents also governs memory allocation in the brain. Although manipulating internal and embodied resources in monkeys would be much more challenging than in artificial agents, the geometry of our navigation task provides a natural asymmetry in embodied-resource reliability that allows us to test this principle. Equal tangential and radial target displacements in the ground plane produce larger changes in horizontal than vertical gaze angle (Fig. 5a). Assuming horizontal and vertical gaze angles are observed with equal, fixed Gaussian noise, horizontal gaze enables a more precise estimate of target position than vertical gaze (Fig. 5b; Methods). The resource-allocation principle thus predicts that horizontal gaze should carry more information about horizontal target location, whereas vertical target location should be represented more independently of gaze in internal states.

**Figure 5:**
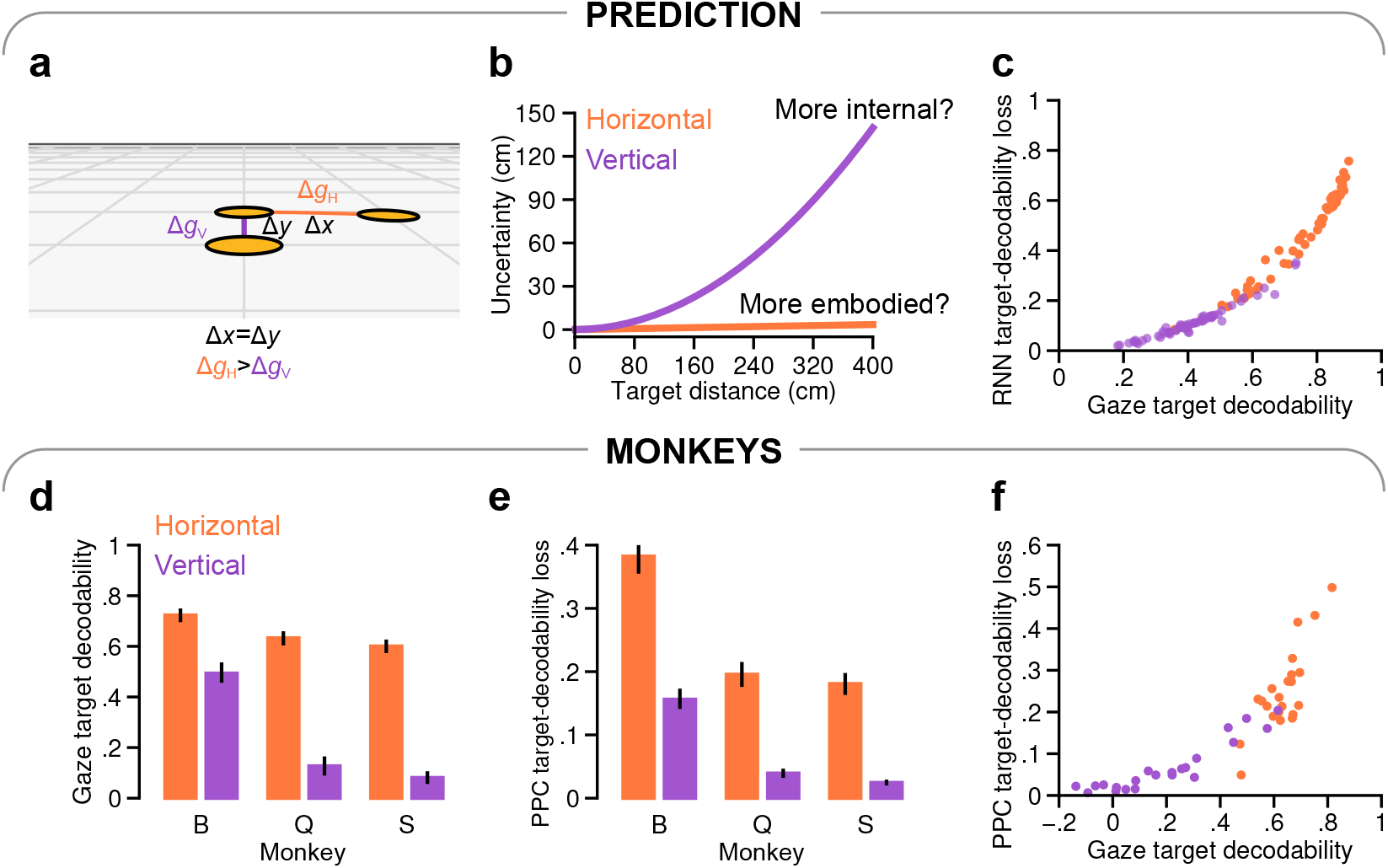
Resource allocation predicts memory organization in macaque parietal cortex. **a**, Task geometry illustrating that equal tangential and radial displacements (Δ*x* = Δ*y*) in the ground plane produce larger changes in horizontal than vertical gaze angle (Δ*g*_*H*_ *>* Δ*g*_*V*_ ). **b**, Predicted target-position uncertainty from horizontal and vertical gaze as a function of target distance, assuming equal Gaussian angular noise (s.d. 0.5°) in the two dimensions and an eye height of 10 cm. Uncertainty was quantified as the inverse square root of Fisher information. **c**, Relationship between gaze target decodability and RNN target-decodability loss across the agent resource landscape, shown separately for horizontal and vertical dimensions. Gaze observation noise had s.d. 0.5 |*g*| for both dimensions, where *g* is gaze angle (Extended Data Fig. 5). **d**, Decodability of hidden-target location from gaze in monkeys B, Q and S for horizontal and vertical dimensions. **e**, Loss of hidden-target decodability from PPC activity after removing gaze-related variance, shown for horizontal and vertical dimensions. **d**,**e**, Error bars indicate 95% CIs across trials. **f**, Relationship between gaze target decodability (**d**) and PPC target-decodability loss (**e**) across recording sessions and dimensions. Pearson correlations were significant overall (*r* = 0.87, *P* = 1.0 ×10^*−*5^) and within horizontal (*r* = 0.82, *P* = 0.004) and vertical (*r* = 0.92, *P* = 2.0 ×10^*−*5^) dimensions (within-animal session-block permutation; Methods). Points denote individual session–dimension pairs. **c–f**, Target decodability was quantified separately for horizontal and vertical dimensions using Frobenius correlation between predicted and true target coordinates. **d–f**, Analyses used 2,000 target-matched trials per subject, sampled across monkey sessions.

We first tested this prediction in agents, where gaze observation noise could be systematically controlled. Agents received signal-dependent gaze observation noise at three levels (Methods). Across the target range, horizontal gaze remained more reliable than vertical gaze, with increasing noise amplifying the difference in target-position uncertainty between dimensions (Extended Data Fig. 5a,b). We then trained agents across the same resource landscape as in Fig. 4 at each observation-noise level. As predicted, increasing observation noise amplified the horizontal–vertical difference in target decodability from gaze and RNN target-decodability loss after gaze removal (Extended Data Fig. 5c–f). Across resource conditions, the two measures remained closely related, while horizontal and vertical dimensions became increasingly separated along this relationship (Fig. 5c; Extended Data Fig. 5g–i). Thus, geometry-dependent differences in gaze reliability predicted corresponding differences in memory allocation across internal and embodied states.

We next tested whether the same prediction held in macaque posterior parietal cortex (PPC), previously shown to encode dynamic estimates of hidden-goal location in the same navigation task [20]. Across all three monkeys, horizontal gaze carried more hidden-target information than vertical gaze (Fig. 5d). To determine how much target information in PPC activity was associated with gaze, we removed gaze-related variance from PPC activity and measured the resulting loss of target decodability (Methods). Neural activity remained highly correlated before and after gaze removal, yet the loss of target decodability was substantially greater in the horizontal than vertical dimension (Fig. 5e; Extended Data Fig. 6a,b). As a control, we shifted gaze in time to disrupt its correspondence with PPC activity, then removed activity variance associated with the shifted gaze signal. Target decodability remained close to its original level, showing that the loss depended on the true gaze–neural relationship (Extended Data Fig. 6b).

Across recording sessions, greater target information in gaze was associated with a greater loss of PPC target decodability after removing gaze-related variance (Fig. 5f). Horizontal target information was generally higher in gaze and showed a greater PPC decodability loss than vertical target information, closely resembling the organization observed in artificial agents (Fig. 5c). Importantly, the association was also present across sessions within the horizontal and vertical dimensions separately, showing that it was not driven solely by the horizontal–vertical separation. Together, these findings reveal a shared principle of memory organization across brains and machines: the relative reliability of internal and embodied resources governs how hidden-goal information is distributed between them.

## Discussion

Our findings identify resource reliability as a determinant of how intelligent systems distribute memory across internal and embodied states. In a naturalistic navigation task, primates and artificial agents maintained information about hidden goals through eye movements, and restricting gaze impaired behavior. Artificial agents revealed that this strategy was not inevitable: gaze became increasingly important as internal memory was degraded, and varying the reliability of internal memory and gaze systematically redistributed goal information between internal and embodied states. The same resource-allocation principle predicted the organization of target representations in macaque parietal cortex: when task geometry made gaze a more reliable resource for a target dimension, more target information was carried by gaze and less was represented independently of gaze in cortical activity. Together, these results suggest that the division of computation across available resources is not fixed, but depends on their reliability.

Embodied accounts of cognition have long proposed that bodily states and environmental interactions can reduce demands on memory, inference and problem solving by changing what must be computed internally [8, 9, 11, 12]. In navigation, we previously found that gaze continues to track latent goals even without optic flow, indicating that this behavior cannot be explained solely by active sensing [18]. What determines whether such an embodied strategy is recruited, however, remained unclear. Our results show that computational constraints can shape the strategy itself: rather than merely limiting performance, they can determine how a problem is solved and where task-relevant information is maintained. This links embodied cognition to broader theories of bounded and resource-limited computation [21, 22, 23, 24].

This perspective has consequences for interpreting neural representations in the context of intelligent behavior. Finding that a task variable is represented in neural activity establishes that the information is present, but does not by itself reveal how that information contributes to the system’s computational strategy. The same task-relevant information may be carried through other resources, while also being reflected in neural activity. Our agent manipulations illustrate this directly: across the resource landscape, similar performance and overall target decodability could mask different allocations of goal information, which became apparent when recurrent activity was considered in relation to gaze. Applying this perspective to macaques then revealed how target information was organized across gaze and gaze-independent parietal representations. Thus, understanding the computational role of a neural representation may require asking not only what information it contains, but how that information relates to the other resources available to support behavior.

Resource availability alone, however, cannot determine whether a bodily or environmental state becomes part of a computation. Even an available and highly reliable resource can be computationally irrelevant if exploiting it does not help solve the task. Its computational value instead depends on whether it preserves, transforms or exposes information in a way that changes the demands placed on internal computation. Our geometric analysis illustrates this dependence: the same oculomotor system provides different amounts of useful information about horizontal and vertical target location because of the structure of the navigation task. Other bodily configurations, actions or environmental interactions may become useful under different task structures because they alter different computational requirements. Thus, understanding resource allocation requires explaining not only how reliable a resource is, but what makes that resource computationally useful for the task.

How resource allocation is acquired and reconfigured remains an open question. In our agents, different resource constraints during training produced different allocations of memory across recurrent and oculomotor states without explicit instruction about how these resources should be used, showing that such organization can emerge through learning. Whether an established allocation can subsequently be reconfigured as resources change remains unresolved: our experiments did not test such reallocation, and resource reliability was not experimentally manipulated in macaques. These manipulations are difficult to perform systematically in biological systems, but artificial agents provide complementary experimental leverage by allowing candidate computational pressures to be varied directly without requiring a shared biological implementation [25, 26]. Here, manipulating internal and embodied reliability revealed a resource-allocation principle that predicted the organization of target information in macaque parietal cortex. This illustrates a complementary experimental loop in which biological observations define phenomena to explain, artificial systems isolate candidate computational principles, and targeted biological experiments test their predictions. The same approach could be used to determine how resource allocation is acquired and reconfigured as resources change.

Taken together, our results identify resource allocation as a principle linking computational constraints to how information is organized across available resources. Embodied memory provides a concrete example of this organization, but the broader implication is that understanding intelligence may require explaining not only what information is represented, but why particular resources become part of the computation in the first place.

## Methods

### Experimental subjects and apparatus

Behavioral and neural recordings were obtained from three adult male rhesus macaques (*Macaca mulatta*; monkeys B, Q and S), aged 7–8 years. The experimental procedures and data collection have been described previously [20]. All experimental procedures were approved by the Institutional Animal Care and Use Committee at Baylor College of Medicine and conducted in accordance with National Institutes of Health guidelines. Monkeys were head-fixed in a primate chair and navigated through a virtual environment using a two-dimensional analog joystick. Visual stimuli were rear-projected onto a screen approximately 30 cm in front of the animals, providing stereoscopic visual feedback.

### Memory-guided navigation task

Monkeys performed a memory-guided navigation task in a virtual environment containing a textured ground plane that provided optic-flow information during movement. The ground texture consisted of small, randomly positioned triangles that were continuously repositioned and reoriented after brief lifetimes (approximately 250 ms), preventing their use as stable spatial landmarks. At the beginning of each trial, a circular target (20 cm radius) appeared at a randomly sampled location within ± 35^*°*^ of the initial heading and at a distance of 1–4 m. The target remained visible for 300 ms before disappearing. Monkeys controlled their linear and angular velocities using a joystick, with maximum speeds of 2 m s^*−*1^ and 90^*°*^ s^*−*1^, respectively. Linear velocity determined forward displacement along the current heading, whereas angular velocity controlled changes in heading. The joystick remained active throughout target presentation and subsequent navigation, and animals were free to move their eyes. Following target disappearance, monkeys navigated using optic-flow information without further visual access to the target. Trials terminated when the animal stopped moving or when the maximum trial duration was reached (5–7 s, depending on the session). Successful navigation was defined as stopping within 65 cm of the target center, for which monkeys received a liquid reward.

### Behavioral recordings

Visual stimuli were generated using OpenGL and updated at 60 Hz. The virtual camera was positioned 10 cm above the ground plane. The animal’s position, linear and angular velocities, eye position, target location and task-event markers were recorded for offline analysis. Heading was reconstructed by integrating angular velocity. Behavioral signals were acquired at 833.33 Hz using Spike2 software (Cambridge Electronic Design). Eye movements were recorded using scleral search coils in monkeys B and Q and video-based tracking in monkey S. Eye positions were expressed as horizontal and vertical gaze angles relative to the head-centered forward direction.

### Neural recordings and spike sorting

Extracellular neural activity was recorded from PPC area 7a in the left hemisphere of all three monkeys using chronically implanted Utah microelectrode arrays. Monkeys B and Q were implanted with 96-electrode arrays arranged on 10 ×10 grids, whereas monkey S received a 48-electrode array arranged on a 6× 8 grid. Electrodes were 1 mm long and spaced 400 *µ*m apart. Broadband neural signals were acquired at 30 kHz using a Blackrock Microsystems recording system. Spikes were detected and sorted offline using KiloSort [27], followed by manual inspection and refinement of spike clusters using Phy. Recording sessions typically yielded 70–100 sorted neurons. Further details are provided in [20].

### Data preprocessing and selection

Spike timestamps were binned at 6-ms resolution (166.7 Hz), consistent with previous analyses of these recordings [28]. Spike counts were smoothed with a Gaussian filter (s.d. 24 ms). Behavioral signals were smoothed using a first-order Savitzky–Golay filter [29] with a 17-sample window (102 ms). Neural and behavioral signals were then averaged over non-overlapping 17-sample windows, yielding a temporal resolution of 102 ms, approximately matching the 100-ms time step of the artificial agents. All preprocessing was performed separately for each trial.

Analyses were restricted to trials in which monkeys actively navigated toward the target, excluding trials in which they failed to initiate movement or failed to stop, consistent with previous studies using the same task [20, 28]. Additionally, trials were excluded if their maximum linear velocity was below 25% of the maximum permitted linear speed or their final displacement from the starting position was less than 30% of the target distance, removing trials with minimal movement or premature termination potentially associated with strategic aborting [30].

### Dynamical systems formulation

We formalized sensorimotor interaction with the environment as a partially observable dynamical system comprising environmental state 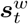, eye state 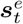nd internal belief *b*_*t*_. Environmental and eye states evolved according to their respective transition dynamics,

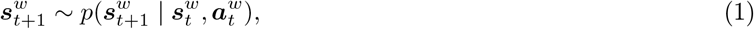

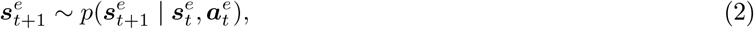

where 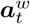and 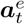denote locomotor and oculomotor actions, respectively. These states generated environmental and eye-state observations,

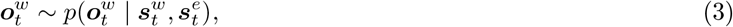

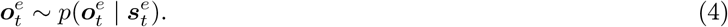

The internal belief was updated from previous belief and current observations,

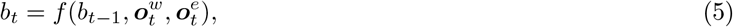

and guided subsequent actions through a policy,

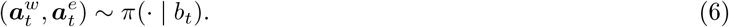

This formulation captures embodied belief formation, in which internal states integrate environmental and bodily information through sensorimotor interaction.

### Modeling the navigation task in artificial agents

Artificial agents were trained in a three-dimensional virtual environment implemented in Unity using the ML-Agents interface [31]. The navigation task followed the macaque experiment, with matching target distributions, locomotor speed limits and reward criteria. Agents were also equipped with an independently controllable eye camera mounted on the navigating body at a height of 20 cm, with a horizontal field of view of 90^*°*^ and a vertical field of view of 60^*°*^. Horizontal and vertical eye movements were controlled through separate velocity commands, with a maximum speed of 150^*°*^ s^*−*1^ and eye positions bounded within ±40^*°*^.

The simulation advanced in 0.1-s steps. At each step, locomotor actions specified linear and angular velocities, which determined forward displacement along the agent’s current heading and changes in heading, respectively. Oculomotor actions independently controlled the horizontal and vertical angular velocities of the eye camera. Actions were bounded within [−1, 1] and scaled by their corresponding maximum velocities. Locomotor commands were additionally corrupted by signal-dependent Gaussian process noise imposed by the environment,

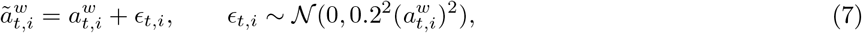

where 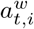denotes a linear or angular locomotor command. No additional process noise was applied to oculomotor commands; their variability arose from stochastic action sampling by the policy (see below).

Each trial began with a target-acquisition period during which agents could move their eyes but could not navigate, allowing them to learn to direct gaze toward visible targets. Target acquisition required the target to fall within a predefined fixation window around the center of the visual field (see below), after which navigation became available. During evaluation, the target disappeared 0.3 s after trial onset, provided it had been acquired. Target disappearance was timed relative to trial onset rather than acquisition, preventing late acquisition from extending target visibility. Navigation began when either locomotor command exceeded 10% of its maximum permitted speed in absolute value and ended when both commands fell below their respective thresholds. Agents received a terminal reward based on their final distance to the hidden target, with no additional rewards for target acquisition or fixation.

### Agent observations

At each time step, agents received nine sensory observations comprising two self-motion signals, two eye-position signals and five visual target-related signals. Self-motion observations represented linear and angular velocities corrupted by signal-dependent Gaussian noise, providing an abstracted representation of optic flow. For each velocity component *v*_*t,i*_, the observation was 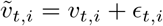, where 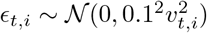. Eye-position observations provided horizontal and vertical angles of the eye camera relative to the navigating body. Independent Gaussian noise was added to each angle, with s.d. *k* |*g*_*t,i*_ |, where *g*_*t,i*_ denotes the corresponding horizontal or vertical eye angle. The eye-position observation-noise coefficient *k* was set to 0.1 by default and varied in the experiments shown in Extended Data Fig. 5. Eye-position noise was applied only after target acquisition.

Visual target observations comprised five signals derived from projecting the target onto the eye camera: the target’s horizontal and vertical screen coordinates, an eccentricity-dependent uncertainty signal, and horizontal and vertical projected sizes. Screen coordinates were normalized to [−1, 1], with zero corresponding to the center of the visual field, whereas projected sizes were expressed as fractions of the camera’s viewport dimensions.

To introduce spatially varying visual uncertainty, observation noise was set to zero within 5^*°*^ of the center of gaze and increased with target eccentricity outside this region, according to

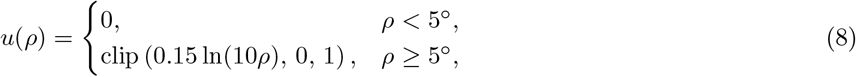

where *ρ* is the target’s eccentricity in degrees, approximated from its normalized screen coordinates scaled by the corresponding horizontal and vertical half-fields of view (45^*°*^ and 30^*°*^, respectively). The uncertainty signal *u*(*ρ*) was provided as the third target observation and also determined the magnitude of Gaussian noise added independently to the screen coordinates and projected sizes. Noise standard deviations were 0.2 |*p*_*i*_ |*u*(*ρ*) for each screen coordinate *p*_*i*_ and 0.1*u*(*ρ*) for each projected size. Following target disappearance, all five target-related observations were replaced by a fixed value of −2.

The observation vector also included the preceding locomotor and oculomotor actions 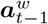and 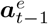, reward *r*_*t−*1_and trial-termination signal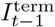, providing efference-copy and task-feedback information. The complete observation vector was therefore

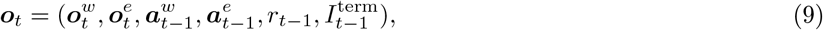

where 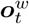 comprises self-motion and visual target observations, 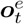 comprises eye-position observations.

### Artificial agent architecture

Artificial agents were implemented in PyTorch using a recurrent actor-critic architecture. The actor contained a gated recurrent unit (GRU) [32] with 128 hidden units, providing an internal state for integrating observations over time. The recurrent output was passed to separate policy heads for locomotor and oculomotor actions. Each head comprised two fully connected layers of 256 units with hyperbolic tangent activations, followed by an output layer specifying the means and standard deviations of two Gaussian action distributions. The critic contained a separate GRU with 256 hidden units, followed by a linear value readout. The actor and critic received the same observation vector but maintained separate recurrent states.

The policy generated four continuous actions comprising linear and angular locomotor commands 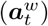 and horizontal and vertical oculomotor commands 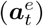. Each action component was sampled independently from a Gaussian distribution,

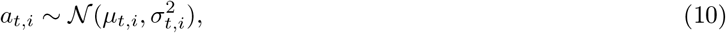

where the action means were bounded by a hyperbolic tangent function and the standard deviations were parameterized as

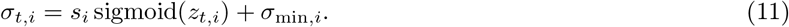

Here, *z*_*t,i*_ denotes the corresponding network output, *s*_*i*_ is a scaling factor and *σ*_min,*i*_ is the minimum action standard deviation. Scaling factors were 0.5 for locomotor actions and 0.3 for oculomotor actions. The minimum standard deviation was fixed at 0.1 for both locomotor dimensions, whereas the minimum oculomotor standard deviation was varied experimentally to manipulate the reliability of embodied actions (see below). Before target acquisition, the minimum oculomotor standard deviation was fixed at 10^*−*5^.

### RL training

Agents were trained using proximal policy optimization (PPO) [33] with generalized advantage estimation (GAE) [34]. The discount factor was *γ* = 0.98, and the GAE parameter was *λ* = 0.98. Policy updates used a clipped surrogate objective with clipping parameter *ϵ* = 0.2, together with a value-function regression loss and an entropy regularization term. The value-loss coefficient was 1 and the entropy coefficient was 0.01.

Network parameters were optimized using Adam [35] with an initial learning rate of 3 × 10^*−*4^ and *ϵ*_Adam_ = 10^*−*3^. Each policy update comprised 80 optimization epochs over 100-step trajectories collected from 500 parallel agents. Rewards were multiplied by a factor of 10 during optimization. Following an initial training stage, the learning rate was reduced to 5 × 10^*−*5^ (see below).

The navigation reward was determined by the Euclidean stopping error *e* between the final stopping position and the target:

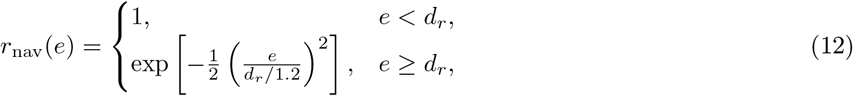

where *d*_*r*_ = 65 cm was the reward boundary. Thus, agents received maximal navigation reward when stopping within the target boundary, with reward decaying exponentially outside the boundary. After the initial training stage (see below), a small penalty proportional to eye eccentricity was also applied at each time step:

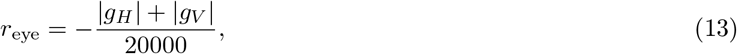

where eye angles were expressed in degrees. No reward was provided for tracking the target with gaze or maintaining target information in oculomotor states.

### Training curriculum

All agents were trained using the same curriculum. During the initial stage, target acquisition was facilitated using a relatively permissive fixation window centered on the visual field, with horizontal and vertical half-widths of 0.1 and 0.05 in normalized screen coordinates, respectively. The target disappearance probability was initially set to 0.8 per step, and no eye-position penalty was applied. The learning rate was 3 × 10^*−*4^.

Once the number of successful trials within a training batch exceeded 200, the fixation tolerances were reduced to 0.05 and 0.025, respectively, and the eye-position penalty was introduced. The per-step target disappearance probability was sampled uniformly between 0.8 and 1 for each trial, and the learning rate was reduced to 5 × 10^*−*5^.

Training was periodically evaluated using a separate Unity environment with the same task parameters as the later training stage, except that the target disappearance probability was fixed at 1. Evaluation used deterministic policy actions and occurred every 25 parameter updates. All agents were trained for 2,000 parameter updates. Different random seeds were used to initialize network parameters and generate training experiences.

### Agent evaluation and checkpoint selection

Agent checkpoints were selected based on navigation performance measured during periodic training evaluations. Performance was quantified as the reward rate during active navigation, defined as the number of rewarded trials divided by cumulative navigation time in seconds, excluding the initial target-acquisition period. For behavioral comparisons (Figs. 1, 2a–d and 3d–f), we selected the first checkpoint at which the reward rate exceeded 0.5 rewarded trials per second, avoiding later training stages at which agent performance could substantially exceed that of monkeys. For all other analyses, we selected the last checkpoint exceeding this threshold, favoring later training stages while maintaining reliable navigation performance. If no checkpoint exceeded the threshold, the checkpoint with the highest reward rate was selected. Checkpoint selection was performed independently for each training seed. All analyses used stochastic actions sampled from the learned policy distributions, except for resource-allocation analyses, which used deterministic actions corresponding to the policy means to isolate differences in learned strategies from differences in action-sampling variability.

### Gaze restriction and perturbation experiments

To assess the functional contribution of gaze to navigation, we restricted oculomotor updating during navigation (Figs. 2f, 3b and 4g). At each time step following target acquisition, horizontal and vertical eye-position updates were blocked with probability *p*, while locomotor control remained unaffected. For Figs. 2f and 3b, agents were trained with *p* = 1, completely preventing eye movements during navigation. For Fig. 4g, gaze restriction was applied only during evaluation, with *p* = 0.8, allowing occasional eye-position updates.

To test the causal influence of oculomotor state on navigation, we introduced additive perturbations to horizontal or vertical oculomotor commands during evaluation (Fig. 3g,h). Perturbations were applied for 10 consecutive time steps (1 s) beginning at navigation onset, with the perturbed action given by

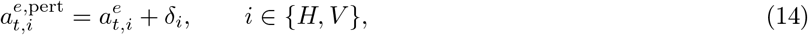

where *δ*_*i*_ denotes the perturbation magnitude. Perturbed commands were converted to eye angular velocities using the maximum oculomotor speed of 150^*°*^ s^*−*1^. Horizontal perturbations ranged from −0.5 to 0.5 in increments of 0.1, whereas vertical perturbations ranged from −0.05 to 0.05 in increments of 0.01. Locomotor actions remained unchanged.

### Internal and embodied resource manipulations

We independently manipulated the reliability of internal and embodied resources during agent training. Internal-memory reliability was controlled by injecting independent Gaussian noise into the actor’s recurrent state at each time step. Specifically, the GRU first computed its updated hidden state

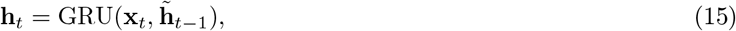

after which Gaussian noise was added to the state propagated to the next time step:

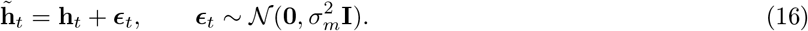

The unperturbed state **h**_*t*_ was used to generate the current action, whereas the perturbed state 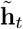was used for subsequent recurrent updates. Increasing *σ*_*m*_ therefore reduced the reliability of information maintained across time steps without directly perturbing the current action output. Noise was applied to the actor’s recurrent state during both training and evaluation, leaving the critic unperturbed to maintain a common value-learning architecture across resource conditions.

Embodied-resource reliability was manipulated by varying the minimum standard deviation *σ*_*e*_ shared by the horizontal and vertical oculomotor action distributions, corresponding to *σ*_min,*i*_ for these two components in the policy parameterization above. This imposed a lower bound on oculomotor action variability during training. Larger values introduced greater irreducible variability in sampled eye commands, whereas smaller values permitted more precise control.

For the initial comparisons of monkey and agent behavior (Figs. 1 and 2a–d), agents were trained with *σ*_*m*_ = 0.2 and *σ*_*e*_ = 10^*−*5^. For the gaze-restriction experiment (Fig. 2f), *σ*_*m*_ was increased to 0.6, while *σ*_*e*_ remained unchanged. To examine how internal-memory constraints influence reliance on gaze (Fig. 3b), we varied *σ*_*m*_ from 0 to 0.8 in increments of 0.1, while holding *σ*_*e*_ = 10^*−*5^. For comparisons between contrasting resource configurations (Fig. 3c–h), embodied-favorable agents were trained with (*σ*_*m*_, *σ*_*e*_) = (0.2, 10^*−*5^), whereas internal-favorable agents were trained with (*σ*_*m*_, *σ*_*e*_) = (0, 0.2).

To systematically characterize memory allocation (Figs. 4 and 5), we trained agents across eight levels of internalmemory noise,

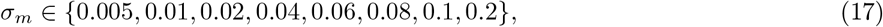

and eight levels of minimum oculomotor action standard deviation,

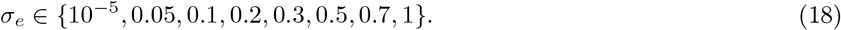

The resulting 8× 8 combinations defined 64 resource configurations. All other network and training parameters were held constant across resource conditions.

### Gaze observation noise

To examine how geometry-dependent differences in embodied reliability influence memory allocation, we additionally manipulated noise in the agent’s eye-position observations (Fig. 5c; Extended Data Fig. 5). Following target acquisition, horizontal and vertical eye positions were observed with independent signal-dependent Gaussian noise,

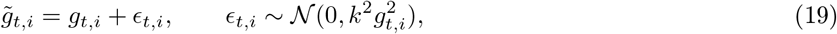

where *g*_*t,i*_ denotes the true gaze angle in dimension *i* ∈ {*H, V}* and *k* controls observation noise. We considered *k* = 0.1, 0.3 and 0.5, applying the same value in both dimensions. At each observation-noise level, agents were trained across the 64 internal- and embodied-resource configurations described above.

### ROC analysis

To quantify navigation accuracy, we performed receiver operating characteristic (ROC) analysis using the Euclidean distance between final stopping locations and target locations. For each trial, stopping distance was calculated relative to the true target and a target randomly shuffled across trials. ROC curves quantified the ability of stopping distance to distinguish true from shuffled target–response pairings, with smaller distances indicating true pairings. Performance was summarized by the area under the ROC curve (AUC).

### Angular coordinates of target and stopping locations

To compare gaze with spatial variables, target and final stopping locations were transformed from world coordinates into angular coordinates relative to the subject’s instantaneous position and heading. For a reference location (*x*_*g*_, *y*_*g*_) and subject position (*x*_*t*_, *y*_*t*_) with heading *ϕ*_*t*_ measured counterclockwise from the positive *x*-axis, the horizontal and vertical angles were computed as

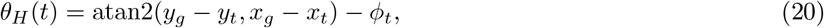

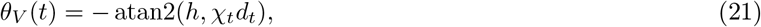

where *h* is eye height, *d*_*t*_ is the Euclidean distance between the subject and the reference location, and *χ*_*t*_∈ {− 1, 1 }indicates whether the reference location lies ahead of (+1) or behind (−1) the subject, based on its projection onto the current heading direction. Angles were expressed in degrees, and horizontal angles were unwrapped to maintain temporal continuity. The same transformation was applied to target and final stopping locations, each treated as a fixed reference in world coordinates. Angular coordinates were excluded once the subject came within the reward boundary of the corresponding reference location. These coordinates were used in most behavioral and neural analyses involving target or stopping locations, except for the steering-error prediction analysis in Fig. 2c (see below).

### Frobenius correlation

Correlations between multidimensional variables were quantified using Frobenius correlation,

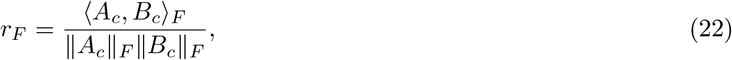

where *A*_*c*_ and *B*_*c*_ are transformed data matrices, ⟨·,·⟩ _*F*_ denotes the Frobenius inner product and ∥ · ∥_*F*_ denotes the Frobenius norm. Each column was transformed according to

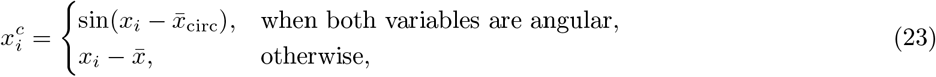

where 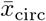and 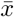 denote the circular and arithmetic means, respectively. For one-dimensional angular variables, this reduces to the Jammalamadaka–Sengupta circular correlation [36]. Unless otherwise specified, correlations throughout the study were computed using this method.

### Trajectory similarity and gaze tracking

To quantify the similarity of steering and gaze trajectories between monkeys and artificial agents, as well as between monkeys (Fig. 1j,l; Extended Data Fig. 1b,c), we compared 2,000 target-matched trial pairs. Trajectories were interpolated to 50 equally spaced samples using two alignment methods: normalized time and normalized movement progress. For time-based alignment, trajectories were interpolated uniformly over their respective durations. For progress-based alignment, trajectories were interpolated according to radial displacement from the starting position, normalized independently for each trial. Trials shorter than 0.5 s were excluded. Trajectory similarity between subject pairs was quantified using Frobenius correlation. Null correlations were obtained by circularly shifting one trajectory within each matched pair by a randomly selected nonzero offset (20 repetitions per trial), with horizontal and vertical dimensions shifted together. Correlations were computed for each trial pair and averaged across pairs.

For visualization in Fig. 1k and Extended Data Fig. 2a,b, vertical gaze, target and stopping-location angles in agents were rescaled to account for differences in eye height between monkeys and agents, assuming gaze intersection with the ground plane. This transformation was not applied to quantitative analyses.

Gaze tracking was quantified by computing Frobenius correlations between gaze angles and the corresponding angular coordinates of the target or final stopping location at each time point following target disappearance. Chance correlations were obtained by shuffling gaze trajectories across trials independently at each time point.

### Predicting navigation errors from gaze

To assess whether gaze target-tracking errors predicted subsequent navigation errors beyond target geometry (Figs. 2d and 3f; Extended Data Fig. 2e), we fitted ridge regression models to predict final linear and angular steering errors. Linear steering error was defined as the Euclidean distance between the target and final stopping location, whereas angular steering error was defined as the signed angular difference between these locations relative to the starting position. Gaze target-tracking error was quantified as the mean Euclidean distance between gaze and target locations in angular coordinates for predicting linear steering error, and as the mean signed horizontal gaze–target difference for predicting angular steering error.

For each outcome, a baseline model used the target’s world coordinates as predictors, whereas a full model additionally included the corresponding gaze tracking error. Models were evaluated using five-fold cross-validation with shuffled trial assignments. Ridge regularization strength was selected using RidgeCV from 60 logarithmically spaced values between 10^*−*4^ and 10^8^. Predictive performance was quantified by the coefficient of determination (*R*^2^) calculated from pooled out-of-fold predictions. The contribution of gaze was defined as 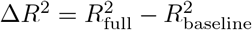.

### Gaze-related variance removal

To quantify gaze-independent components of population activity, we removed gaze-related variance using multivariate linear regression. Population activity comprised recurrent hidden states in artificial agents or PPC neural activity in monkeys. For population activity **x**_*t*_ and horizontal and vertical gaze angles **g**_*t*_, we fitted

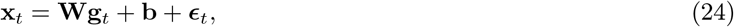

and obtained gaze-removed activity as

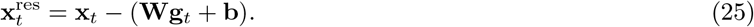

Regression coefficients were estimated using all available time points within each agent dataset or monkey recording session. As a control, gaze trajectories were circularly shifted by a randomly selected nonzero offset independently within each trial, with horizontal and vertical gaze shifted together. The same regression procedure was repeated using shifted gaze to obtain control residual activity.

### Target decoding

Hidden-target location, expressed in angular coordinates, was decoded from gaze, original population activity and gaze-removed population activity and shifted-gaze control residual activity using ridge regression. Horizontal and vertical target coordinates were decoded separately using five-fold GroupKFold cross-validation, with entire trials assigned to the same fold. Decoding was performed separately for each agent seed and each monkey recording session. Regularization strengths were selected using RidgeCV from 60 logarithmically spaced values between 10^*−*4^ and 10^8^, independently for horizontal and vertical target coordinates. Decodability was quantified by Frobenius correlation between out-of-fold predictions and true target coordinates. Target-decodability loss following gaze removal was defined as the difference between decodability from original and gaze-removed population activity.

### Spatial information under constant angular gaze noise

We quantified how ground-plane geometry constrains the spatial information available from horizontal and vertical gaze position (Fig. 5b). We assumed that a gaze angle *g*(*x*) encodes a spatial variable *x* with additive Gaussian noise of constant variance 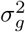. The Fisher information about *x* is

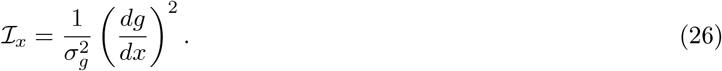

We defined the Fisher-information-limited spatial uncertainty as 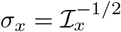.

For gaze directed toward a target on the ground plane at distance *d* from an observer with eye height *h*, the magnitude of the vertical eye angle is

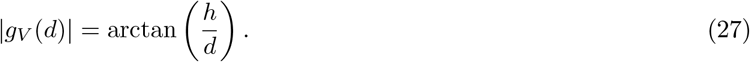

The Fisher information about radial distance is therefore

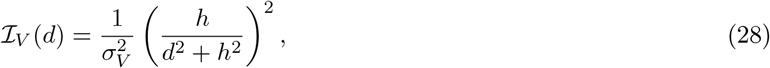

with corresponding spatial uncertainty

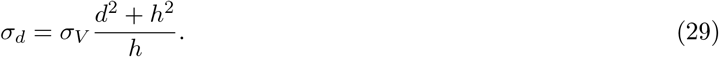

For *d* ≫ *h*, these expressions reduce to

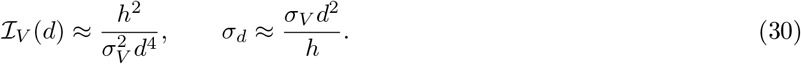

For horizontal eye position, an infinitesimal tangential displacement *dx*_*⊥*_ at distance *d* produces an angular change of magnitude |*dg*_*H*_ | = |*dx*_*⊥*_|*/d*. The corresponding Fisher information about tangential position is

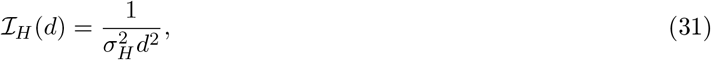

with spatial uncertainty

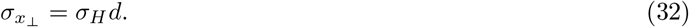

Thus, under constant angular gaze noise, horizontal spatial uncertainty increases linearly with viewing distance, whereas vertical spatial uncertainty increases quadratically for *d* ≫ *h*.

### Spatial information under gaze-dependent angular noise

Because the magnitude and form of gaze-position noise in biological systems are not known, we additionally considered a scale-free noise model in which angular noise was proportional to gaze eccentricity (Extended Data Fig. 5), matching the multiplicative observation noise used in artificial agents:

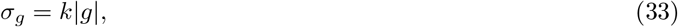

where *k* is a dimensionless relative noise coefficient.

As in the constant-noise analysis, we quantified spatial information from the sensitivity of mean gaze angle to spatial position, with local spatial uncertainty defined as the inverse square root of this information. Under relative noise, this corresponds to the mean-sensitivity component of Fisher information, excluding the additional contribution from parameter-dependent observation variance.

For vertical gaze position, substituting *σ*_*V*_ = *k*|*g*_*V*_ | = *k* arctan(*h/d*) gives

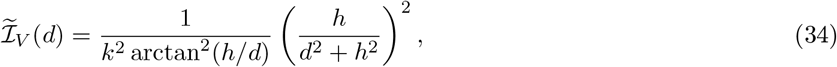

with corresponding spatial uncertainty

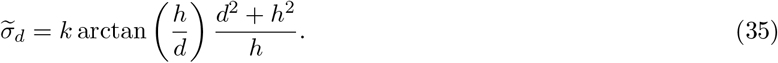

For *d* ≫ *h*, these expressions reduce to

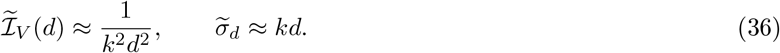

For horizontal gaze position, we considered infinitesimal tangential displacements perpendicular to the target direction at radial distance *d*. Relative angular uncertainty depends on horizontal gaze eccentricity *g*_*H*_ . For nonzero eccentricity, the information about tangential position is

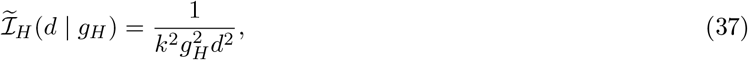

with corresponding spatial uncertainty

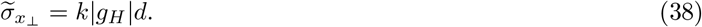

Thus, under relative angular noise, uncertainty in both radial and tangential position increases approximately linearly with viewing distance for *d* ≫ *h*. Their relative magnitudes depend on horizontal gaze eccentricity:

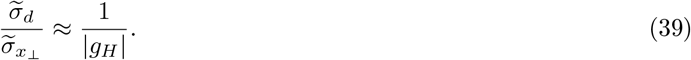

Because the maximum target eccentricity in our task was less than 1 rad, the model predicts lower horizontal than vertical spatial uncertainty for target-directed gaze in this limit.

## Statistical analysis

Statistical analyses were performed in Python. Throughout the study, correlation coefficients were averaged in Fisher-*z* space and transformed back for reporting. Unless otherwise indicated, 95% CIs were defined by the 2.5th and 97.5th percentiles of the bootstrap distribution.

For trajectory similarity analysis (Fig. 1j,l), correlations were first averaged across target-matched trial pairs within each recording session. For agent–monkey comparisons, session-level correlations were additionally averaged across independently trained agent seeds. Mean correlations and 95% CIs were then calculated separately for each monkey by resampling recording sessions with replacement 5,000 times.

For time-resolved gaze correlations (Figs. 2b and 3e), 95% CIs were estimated using 1,000 paired bootstrap resamples of trials at each time point. Differences between correlations were evaluated using paired bootstrap resampling. For shuffled controls, gaze observations were circularly shifted along the trial dimension at each time point, using 20 randomly sampled nonzero offsets.

For comparisons of predictive performance (Figs. 2d and 3f), 95% CIs were estimated using 100,000 paired bootstrap resamples of trials. Observed outcomes and corresponding out-of-fold predictions from the baseline and full models were resampled together, and differences in predictive performance were calculated within each bootstrap sample.

For monkey decoding analyses (Fig. 5d,e), 95% CIs were estimated using 10,000 bootstrap resamples of entire trials. Differences between decoding correlations were evaluated using paired trial-level bootstrap resampling. For Extended Data Fig. 5d,f, 95% CIs were estimated by bootstrapping resource conditions.

For correlations between gaze decodability and PPC target-decodability loss (Fig. 5f), significance was assessed using 100,000 one-sided permutations. PPC decodability-loss values were permuted across sessions within each monkey, preserving paired horizontal and vertical measurements. Pearson correlations were recomputed for pooled and individual dimensions. *P* values were calculated as (*N*_extreme_ + 1)*/*(*N*_perm_ + 1), where *N*_extreme_ is the number of permuted correlations at least as large as the observed correlation. For other Pearson correlation tests, *P* values were obtained using SciPy’s default two-sided test.

## Data availability

The data supporting the findings of this study are available at https://github.com/ryzhang1/embodied_memory.

## Code availability

The code and instructions required to reproduce the analyses and figures are available at https://github.com/ryzhang1/embodied_memory.

## Acknowledgments

We thank Jing Lin and Jian Chen for assistance with stimulus programming; Yizhou Chen for useful discussions; Kaushik Lakshminarasimhan, Eric Avila, and Erin Neyhart for data collection; Karen Wood and Rebecca Meyer for spike sorting; Roozbeh Kiani for surgical expertise; and Antara Ghosh for assistance with scientific illustrations. This work was supported by NIH CRCNS grant 1R01 NS120407-01 and Simons Foundation SCENE grant NC-SCN-00007276-10 (X.P. and D.E.A).

## Author contributions

Conceptualization and methodology: R.Z., X.P., and D.E.A. Investigation and formal analysis: R.Z. Visualization: R.Z. Supervision: X.P. and D.E.A. Writing—original draft: R.Z. Writing—review and editing: R.Z., X.P., and D.E.A. Funding acquisition: X.P. and D.E.A.

## Extended data figures and tables

**Extended Data Fig. 1:**
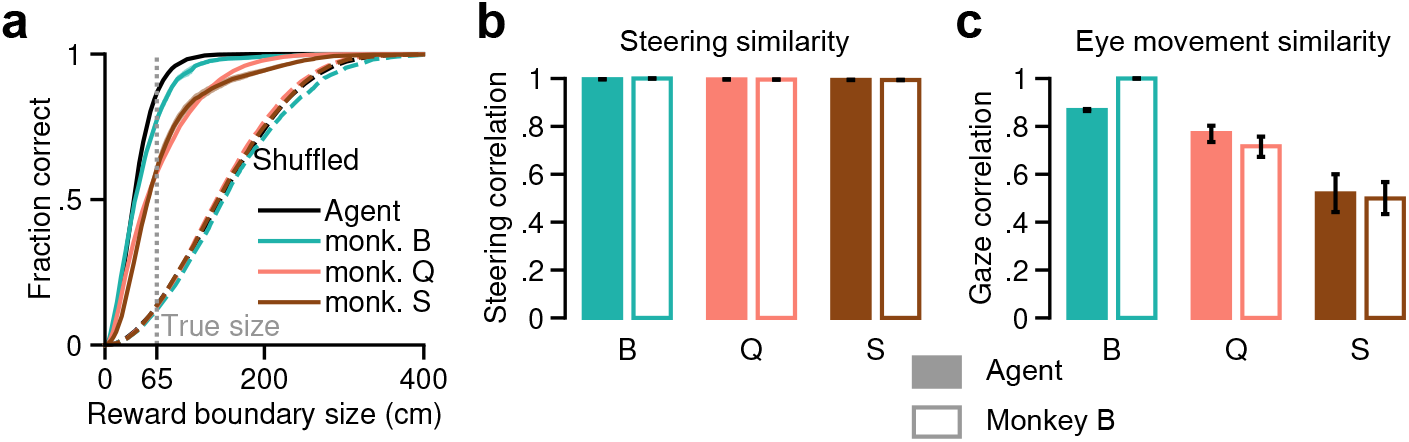
Quantification and robustness of navigation and gaze behavior. **a**, Fraction of correct trials as a function of hypothetical reward-boundary size. Solid lines denote true target–response pairings; dashed lines denote pairings after shuffling targets across trials. Grey dotted line indicates the actual reward boundary. Analyses used 2,000 trials per subject. **b**,**c**, As in Fig. 1j,l, respectively, but trajectories were aligned by navigation progress, defined as Euclidean displacement from the trial starting position, rather than elapsed time.

**Extended Data Fig. 2:**
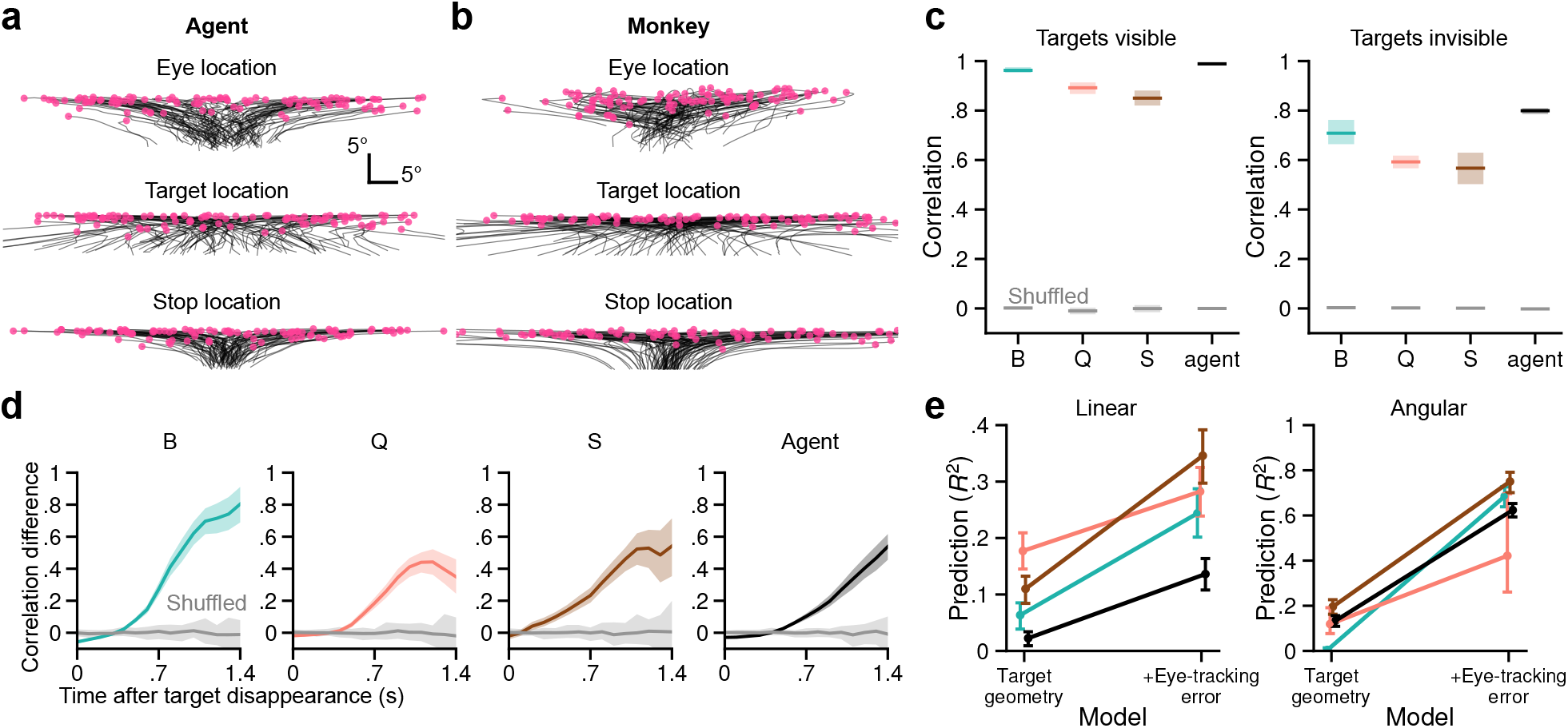
Relationship between gaze, hidden goals and subsequent navigation. **a**,**b**, Example eye, target and final stopping locations in angular coordinates across trials for an agent (a) and monkey B (b). **c**, Frobenius correlations between gaze and target location when targets were visible (left) or invisible (right), collapsed across time. Grey indicates chance correlations obtained by independently shuffling targets across trials; shading indicates 95% CI. **d**, Difference between gaze–final-stop and gaze–target correlations over time after target disappearance. Grey indicates chance obtained by shuffling gaze across trials; shading indicates 95% CI. **e**, Cross-validated out-of-sample *R*^2^ for predicting final linear (left) and angular (right) steering errors using target geometry alone or together with gaze target-tracking error. Error bars indicate 95% CI. **c–e**, Analyses used 2,000 target-matched trials per subject, sampled across monkey sessions or trained agent seeds.

**Extended Data Fig. 3:**
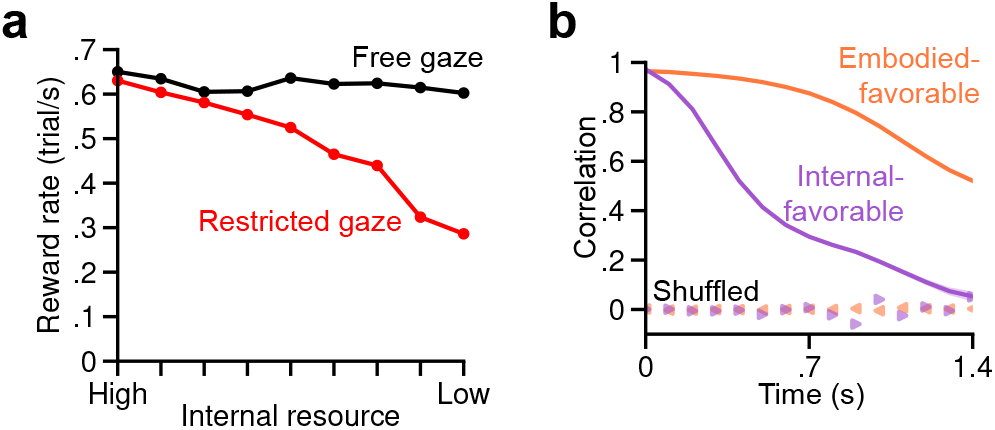
Additional analyses of internal-resource constraints. **a**, As in Fig. 3b, but showing reward rate, defined as rewarded trials per second. **b**, As in Fig. 3e, but showing gaze–final-stop correlation.

**Extended Data Fig. 4:**
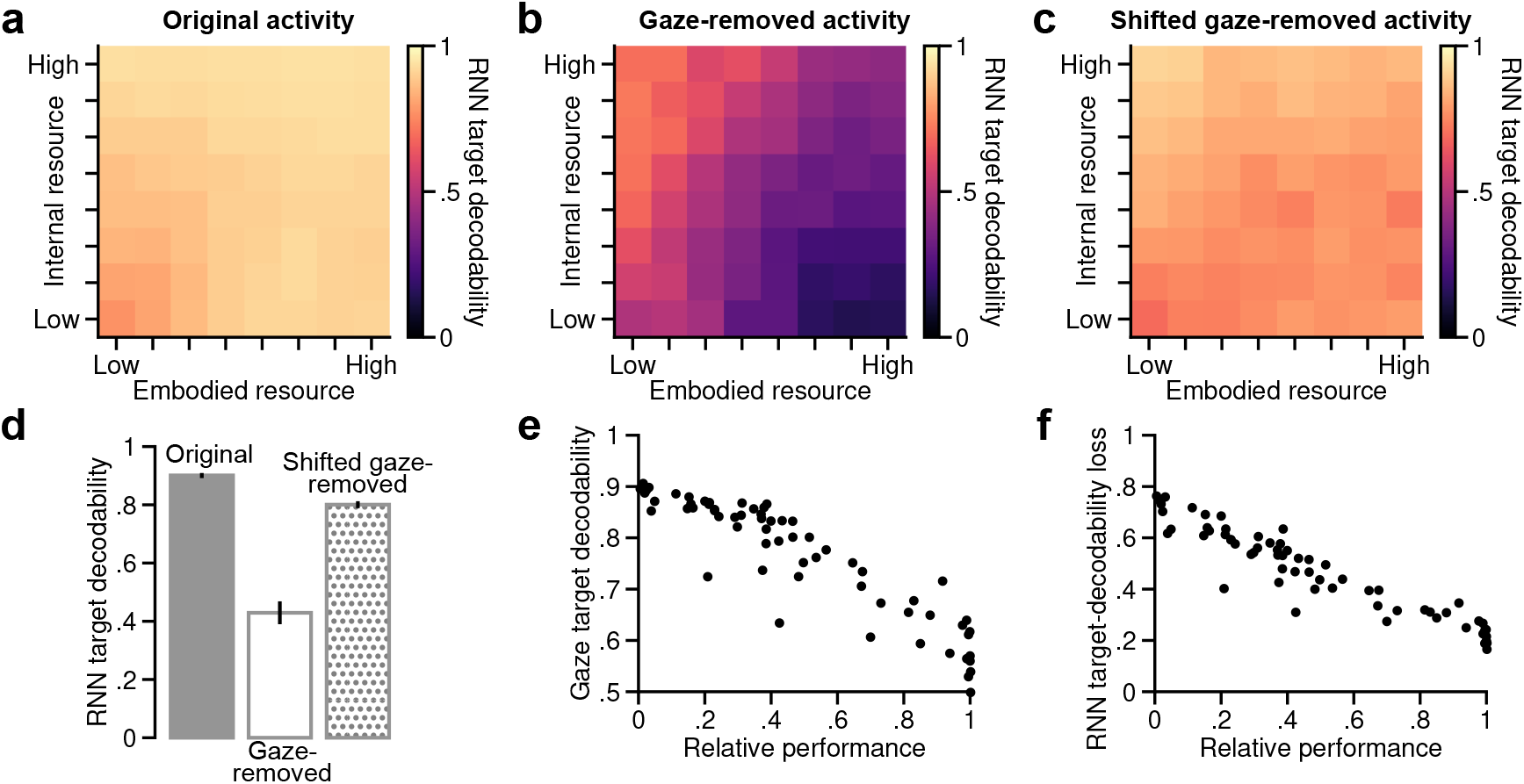
Controls and behavioral relevance of resource-dependent target representations. **a–c**, Decodability of hidden-target location from original recurrent activity (**a**), gaze-removed activity (**b**) and shifted gaze-removed activity (**c**) across the resource landscape. For the shifted-gaze control, gaze was circularly shifted within each trial before gaze-related variance was removed from recurrent activity. Decodability was quantified by Frobenius correlation between predicted and true target locations. Values are averaged across eight training seeds per resource condition. **d**, Target decodability from **a–c** averaged across the 64 resource conditions. Error bars indicate 95% CI. **e**, Relationship between gaze target decodability (Fig. 4d) and relative performance under gaze restriction (Fig. 4g) across the 64 resource conditions (Pearson’s *r* = −0.92, *P* = 6.3 ×10^*−*27^). **f**, Relationship between recurrent target-decodability loss (Fig. 4e) and relative performance under gaze restriction (Fig. 4g) across the 64 resource conditions (Pearson’s *r* = −0.95, *P* = 2.6 × 10^*−*32^).

**Extended Data Fig. 5:**
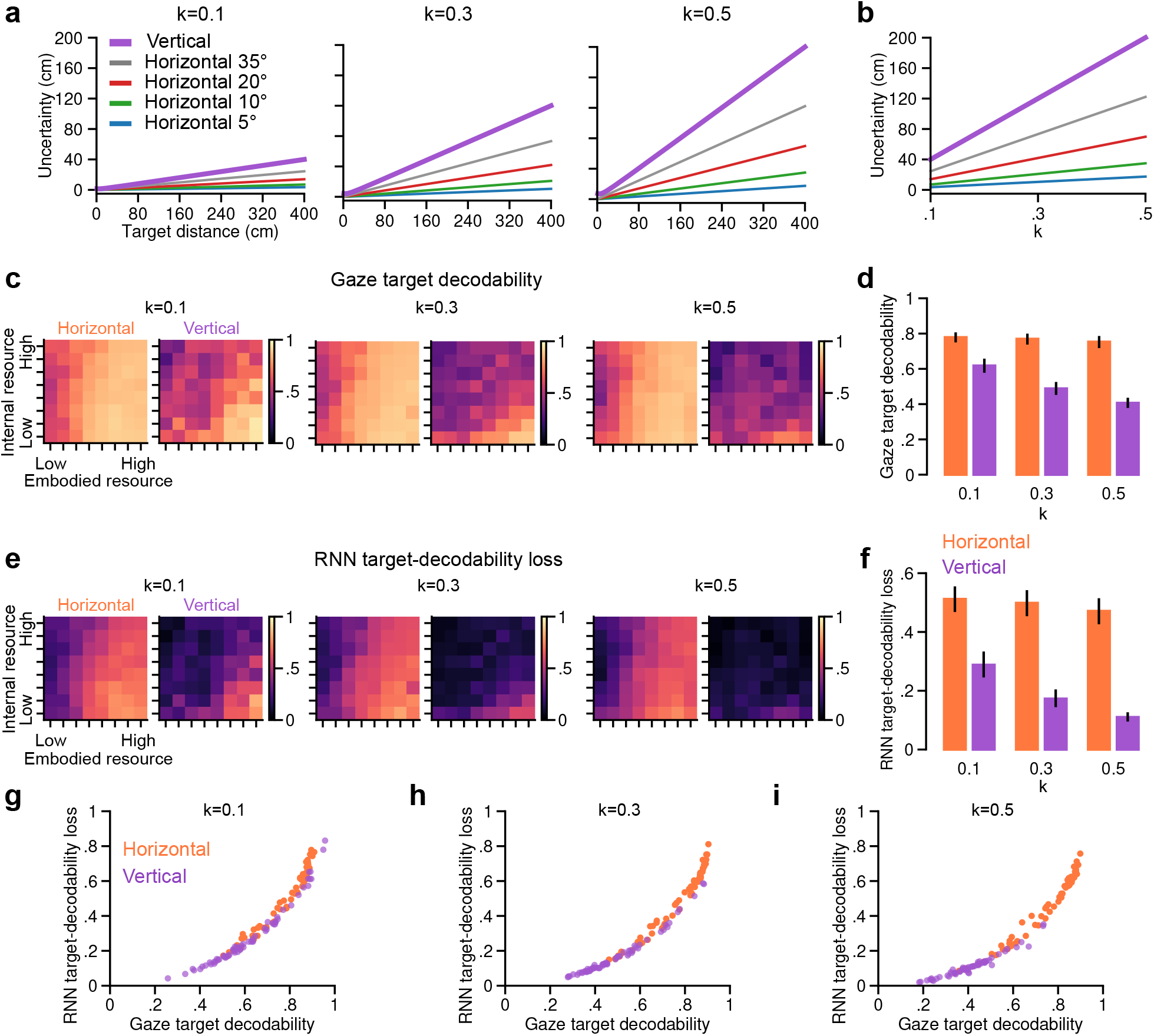
Gaze observation noise modulates geometry-dependent memory allocation. **a**, Predicted target-position uncertainty from horizontal and vertical gaze as a function of target distance. Gaze observation noise was Gaussian with s.d. *k* |*g*|, where *g* is gaze angle and *k* = 0.1, 0.3 or 0.5, with equal *k* in the two dimensions. Horizontal uncertainty is shown for horizontal target angles of 5°, 10°, 20° and 35°; vertical uncertainty is angle-independent. **b**, Predicted target-position uncertainty as a function of *k* at a target distance of 400 cm for vertical gaze and the horizontal gaze angles shown in **a. c**, Gaze target decodability across the resource landscape for *k* = 0.1, 0.3 and 0.5, shown separately for horizontal and vertical dimensions. **d**, Gaze target decodability for each *k* and dimension in **c**, averaged across resource conditions. **e**, As in **c**, but showing RNN target-decodability loss after gaze removal. **f**, RNN target-decodability loss for each *k* and dimension in **e**, averaged across resource conditions. **d**,**f**, Error bars indicate 95% CI across resource conditions. **g–i**, Relationship between gaze target decodability and RNN target-decodability loss across the agent resource landscape for horizontal and vertical dimensions at *k* = 0.1 (**g**), 0.3 (**h**) and 0.5 (**i**). The *k* = 0.5 condition is shown in Fig. 5c. **c–i**, Values are averaged across eight training seeds per resource condition. Target decodability was quantified separately for horizontal and vertical dimensions using Frobenius correlation between predicted and true target coordinates.

**Extended Data Fig. 6:**
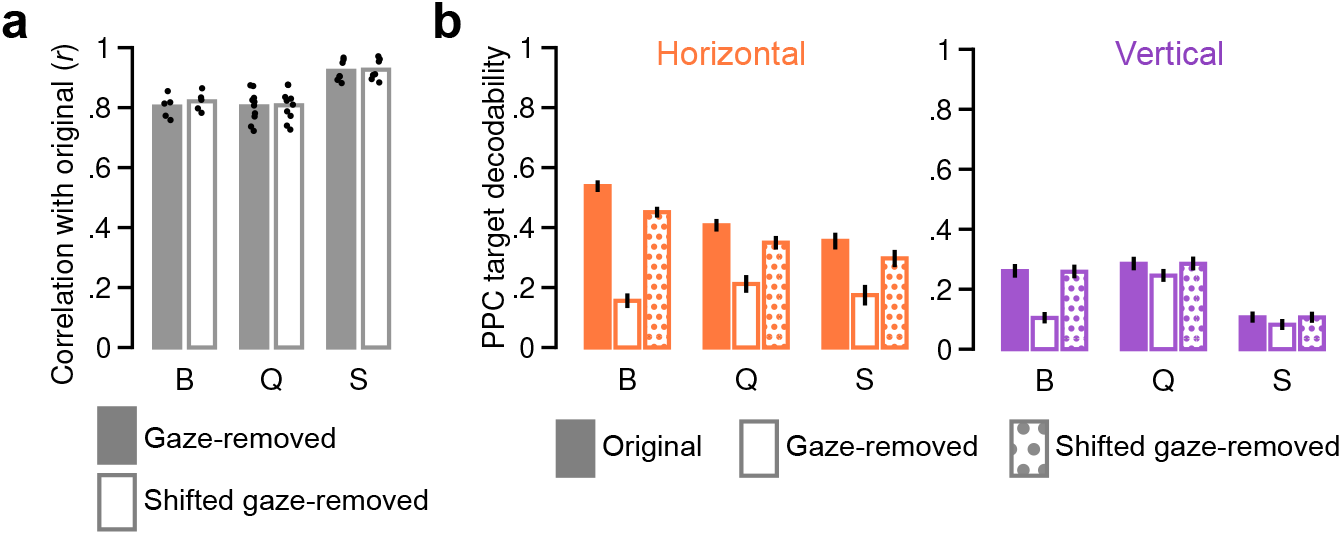
Gaze-removal controls and target decodability in macaque parietal cortex. **a**, Pearson correlation between original PPC activity and gaze-removed (filled) or shifted gaze-removed (open) activity in monkeys B, Q and S. Bars indicate means across recording sessions; black dots denote individual sessions. **b**, Decodability of hidden-target location from original, gaze-removed and shifted gaze-removed PPC activity, quantified separately for horizontal (left) and vertical (right) coordinates using Frobenius correlation between predicted and true values. Error bars indicate 95% bootstrap CIs across trials. **a**,**b**, Gaze-related variance was linearly removed from PPC activity; for the shifted-gaze control, gaze was circularly shifted within each trial before residualization. Analyses used 2,000 target-matched trials per subject, sampled across monkey sessions.

